# Stem cell delivery to kidney *via* minimally invasive ultrasound-guided renal artery injection in mice

**DOI:** 10.1101/609230

**Authors:** May Zaw Thin, Olumide Ogunldade, Joan Comenge, P. Stephen Patrick, Daniel J. Stuckey, Anna L.David, Raphaël Lévy, Mark F. Lythgoe, Paul Beard, Tammy L. Kalber

## Abstract

Cell-based therapies are promising treatments for various kidney diseases. However, the major hurdle in initiating therapeutic responses is the inefficiency of injection routes to deliver cells to the kidney parenchyma. Systemic injection, such as intravenous injection only delivers a small proportion of cells to the kidney. Whereas direct delivery, such as renal artery injection require surgical procedures. A minimally invasive renal artery injection was therefore developed to enhance cell delivery to kidney. In this study, luciferase expressing human adipocyte derived stem cells (ADSC) were labelled with gold nanorods (GNR) and injected into the renal artery using ultrasound guidance. The ADSCs were tracked using bioluminescence and photoacoustic imaging serially over 7 days. Imaging confirmed that the majority of signal was within the kidney, indicative of successful injection and that signal remained stable for 3 days. Histology showed co-localization of GNRs with ADSC staining throughout the kidney with no indication of injury caused by injection. These findings demonstrate that ultrasound-guided renal artery injection is feasible in mice and can successfully deliver a large proportion of cells which are retained within the kidney for 3 days. Therefore, the techniques developed here will be useful for optimising cell therapy in kidney diseases.

## Introduction

Over the past few years, stem cell-based regenerative therapy has been vastly studied as an alternative treatment for various kidney diseases. Mesenchymal stem cells (MSCs) have been used to repair kidney damage due to their ability to release a variety of active biological factors including growth factors such as insulin-like factor 1 (IGF-1)^1^, vascular endothelial growth factor (VEGF)^2^, and other immune system signalling molecules, such as, transforming growth factor-β (TGF-β)^3^. Through these signalling cascades, MSCs are able to recruit and promote repair processes during acute kidney injury (AKI). Therefore, MSC-based therapy has been tested in several AKI mouse models and has been used to treat patients with AKI in three clinical trials^4^.

Although many studies have demonstrated the efficacy of stem cell therapy in kidney diseases, the engraftment of stem cells within the kidney parenchyma as a routine clinical treatment is hindered by the inefficient delivery and retention of stem cells after transplantation. Intravenous (IV), intra-arterial and intraparenchymal injections have been tested in many mouse models of AKI to deliver MSCs to kidney. Although IV injection is a relatively easy and non-invasive method of administration, the main problem is the pulmonary entrapment of cells directly after injection which thereby reduces engraftment within the kidney^5^. Several studies have shown that systemic intra-arterial delivery which by-passes the lung can provide increased cell retention in kidney^6,7^. However, evidence of vascular occlusion after intra-arterial injection has been reported which raises safety concerns^8^. A study by *Alfarano et al.*, directly injected cells into the renal parenchyma which resulted in reduced kidney fibrosis in a rat model of renal ischemia-reperfusion^9^ but this approach is difficult to implement in a clinical setting due to the risk of additional injury. Similarly, the delivery of MSCs directly into the renal artery of an injured kidney has been shown to increase cell engraftment in preclinical studies^10,11^. However, this method is invasive and is usually conducted at the end of the open abdominal surgery to induce renal ischemia. Therefore, the development of a less invasive and more efficient injection route is urgently required to improve stem cell delivery to preclinical kidney injury models.

In addition to establishing the optimal route for stem cell delivery, tracking of transplanted cells and verifying their viability within the target organ is equally important in determining therapeutic efficacy of the cell therapy. Nuclear imaging using ^111^In-oxine labelling has been used to track stem cells in a mouse model of renal ischemia-reperfusion injury^12^. However, due to its low spatial resolution, it lacks detailed information on cell localisation. Iron oxide particles are one of the most widely used MRI-based cell tracking agents for locating transplanted cells in kidney^10,13,14^. Although MRI can provide excellent spatial resolution, iron oxide particles produce negative contrast which makes it difficult to quantify the number of cells in kidney. In addition, the presence of iron oxide particles will interfere with potential functional MRI assessments utilized for kidney diseases such as blood flow measurements.^15^ Despite many studies having explored the applicability of various fluorescent probes, such as near infrared dyes for stem cell tracking in kidney^16,17^, the utility of optical imaging is limited owing to light scattering which reduces the spatial resolution and penetration depth.

Compared to current imaging modalities, photoacoustic (PA) imaging is a relatively new but rapidly expanding non-invasive medical imaging field. It is a combined technology of ultrasound and optical imaging. The mechanism of PA is based on the detection of light absorption in ultrasound (acoustic) wave form generated by laser excited chromophores within tissue, for example, haemoglobin. Since acoustic waves are not scattered as much as light photons, the advantages of PA are deeper tissue penetration and better spatial resolution than optical imaging. A previous study has demonstrated the great potential of PA as a cell tracking tool by using a tyrosinase-based genetic reporter^18^. However, the production of melanin from this reporter gene can be cytotoxic to some cell types and therefore may not be suitable for stem cell tracking^19^. Therefore, to use PA imaging in tracking cell therapies, the contrast agents need to have minimal cytotoxicity with little interference in the cell’s biological functions.

Since gold is known to be non-cytotoxic and inert in biological environment, gold nanoparticles have been extensively used *in vitro* cell imaging^20^. Gold nanoparticles are also promising PA contrast agents due to their unique optical properties which allow optical absorption in the near infrared window where background tissue absorbance is minimal. In addition, the sensitivity of PA imaging can be improved by modifying the shapes of gold nanoparticles such as gold nanorods (GNRs) because of their tunable optical wavelength which results from surface plasmon resonance^21^. Moreover, unlike iron oxide nanoparticles, GNRs do not impede any functional assessment of the target organ by MRI. For these reasons, GNRs have been successfully applied as stem cell tracking agents for PA imaging^22^. As many studies have showed the advantages of PA imaging in detecting kidney disease such as ischemic kidney damage^23^, early kidney injury^24^, Adriamycin-induced nephropathy^25^ and polycystic kidney diseases^26^, tracking GNRs labelled MSCs in kidney with PA imaging will provide accurate cell localisation which corresponds to the structural and functional status of the kidney.

In this study, a novel non-surgical ultrasound-guided renal artery injection was developed to improve stem cell delivery to the kidney without the need for open abdominal surgery. In addition, a dual bioluminescence imaging (BLI) and PA imaging approach was applied in this study by labelling luciferase expressing ADSCs with GNRs. This allowed for the visualisation of cell viability (BLI) with cell localisation within the kidney serially over time after ultrasound-guided renal artery injection.

## Result

### Intracellular uptake of GNRs and their effect on cell proliferation and differentiation potential

Internalisation of GNRs by ADSCs was visualised under a light microscope using sliver enhancement staining which showed the presence of dark aggregates inside the cytoplasm of ADSCs compared to control (Figure **1a** & **b**). The effect on cell proliferation of ADSCs after GNR labelling was studied by measuring bioluminescence emission as a surrogate measurement of cell proliferation. There were no differences in signal intensity between control and GNR labelled groups which showed an increase in signal from day 1 to day 3 after plating which remained stable throughout the study (Figure **1c**). The effect on tri-lineage (adipogenic, chondrogenic and osteogenic) differentiation potential of ADSCs after GNR labelling was assessed by performing differentiation assays (Figure **2**). The results showed both the control and GNR labelled cells could differentiate towards tri-lineages at a similar rate. These results indicate that GNR labelling has no adverse effect on cell proliferation and differentiation potential of ADSCs.

**Figure 1.**
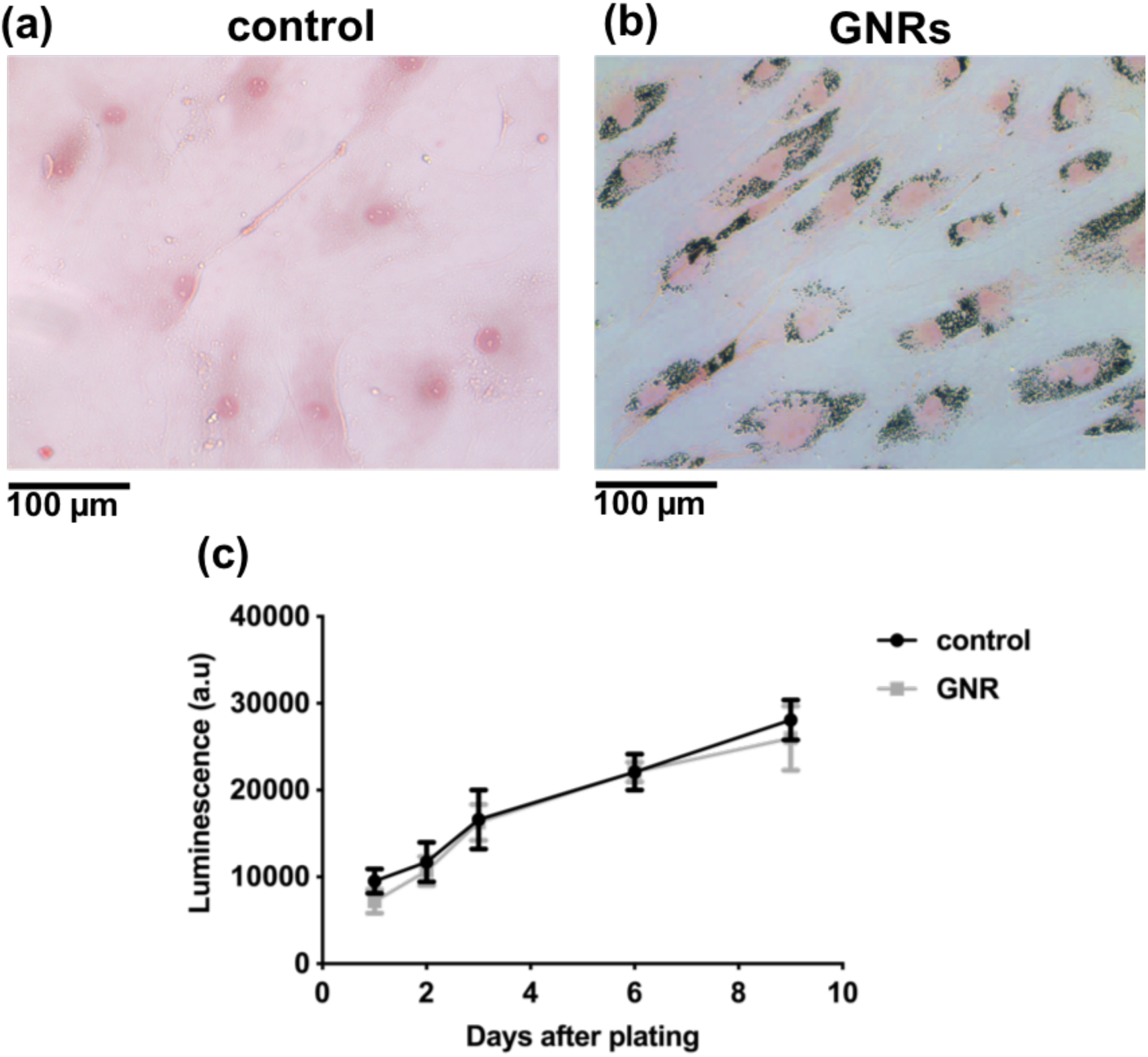
Intracellular uptake of GNRs in ADSCs and the effect on cell proliferation. (a & b) Sliver enhancement staining of ADSCs treated with GNRs showing the dark aggregates inside the cytoplasm compared to untreated control (scale bar = 100 µm). (c) Luciferase-based cell proliferation assay at different time points showing no significant difference between BLI signals (photons/sec in log scale) of control and GNR labelled ADSCs (data are shown as mean ± SD, n = 3).

**Figure 2.**
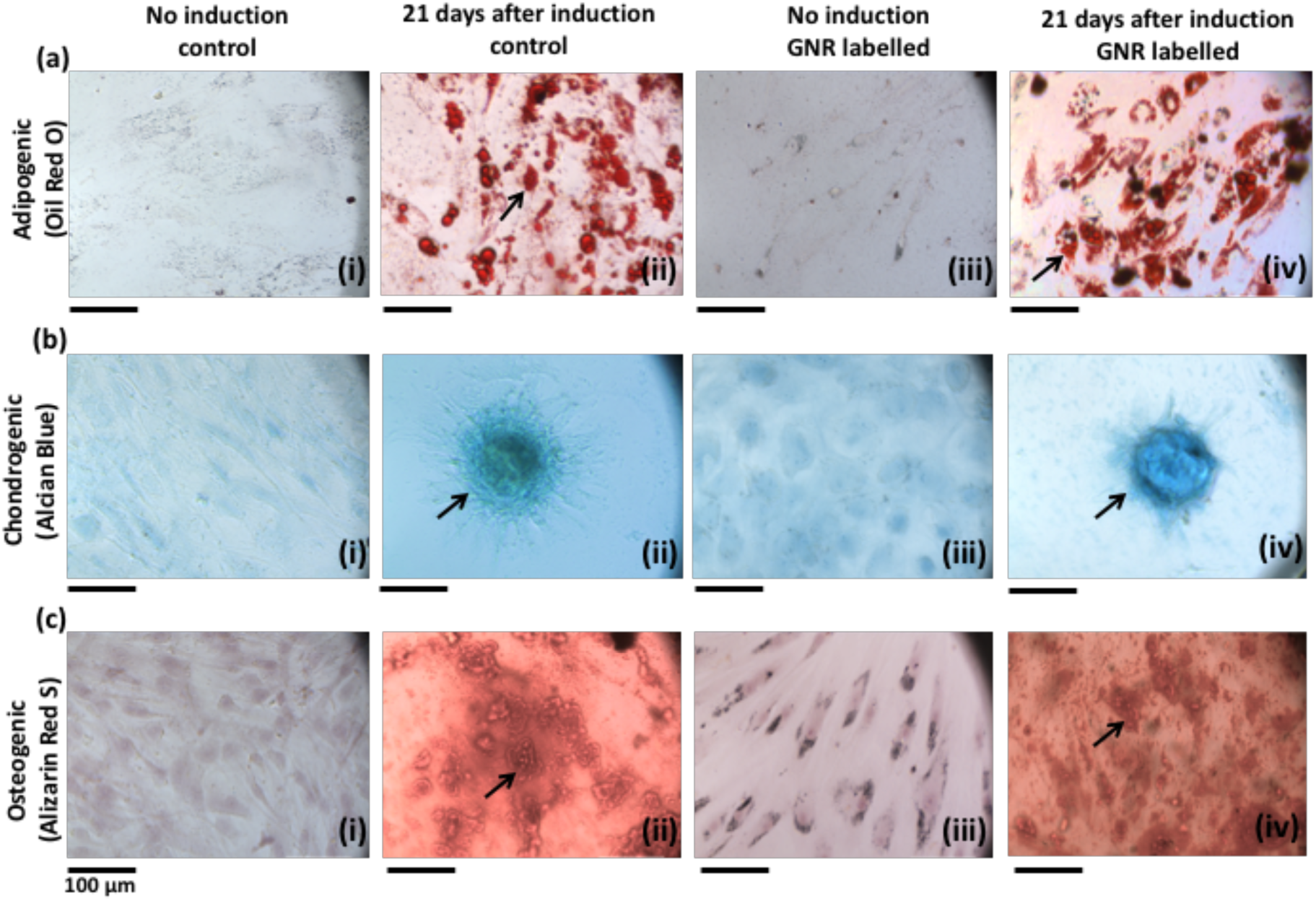
Tri-lineage differentiation of control ADSCs and GNR labelled ADSCs. (a) Oil red o staining for adipogenic differentiation which displays the red coloured oil droplets (indicated by arrows in (ii & iv)). (b) Alcian blue staining for chondrogenic differentiation which displays the blue coloured proteoglycans (indicated by arrows in (ii & iv)). (c) Alizarin red s staining for osteogenic differentiation which displays the red colored calcium deposits (indicated by arrows in (ii & iv)). Scale bar = 100 µm.

### Stem cell delivery to kidney *via* ultrasound-guided renal artery injection

Ultrasound-guided renal artery injection was performed using 6-8-week-old female nude mice. Under anaesthesia, mice were positioned in the left lateral position on the ultrasound platform with the limbs secured with tape. The injection mount with syringe holder was positioned towards the paravertebral muscle of the animal (Figure **3a**). Ultrasound gel was applied to the right lower quadrant of the abdomen and the right kidney was located by using the liver as reference. A colour Doppler image was acquired to visualise the blood vessels of the right kidney (Figure **3b**). In order to identify the arterial and venous flow velocity waveforms of the kidney blood vessels, a pulsed wave Doppler sample gate was placed within the vessels at the appropriate angle relative to flow direction (Figure **3c** & **d**). 2 × 10^5^ GNR labelled ADSCs were suspended in 100 μl of 0.6% alginate solution and injected slowly into the renal artery using a 1 ml syringe with a 29 g needle. In order to avoid surrounding organs damage, the needle was penetrated into the renal artery by passing through paravertebral muscle (Figure **3e**). Successful injection was visualised by the presence of hyperechogenic contrast from the alginate solution around the renal cortex (Figure **3f**). The needle was slowly withdrawn from the body and a colour Doppler image was acquired to assess blood flow to the kidney. The heart rate, body temperature and respiration rate were monitored during the injection and imaging of the kidney which took approximately 15 minutes. The animals were then recovered from anaesthesia and no significant weight loss or sign of major trauma was observed as monitored by body weight and colour of the urine throughout the study.

**Figure 3.**
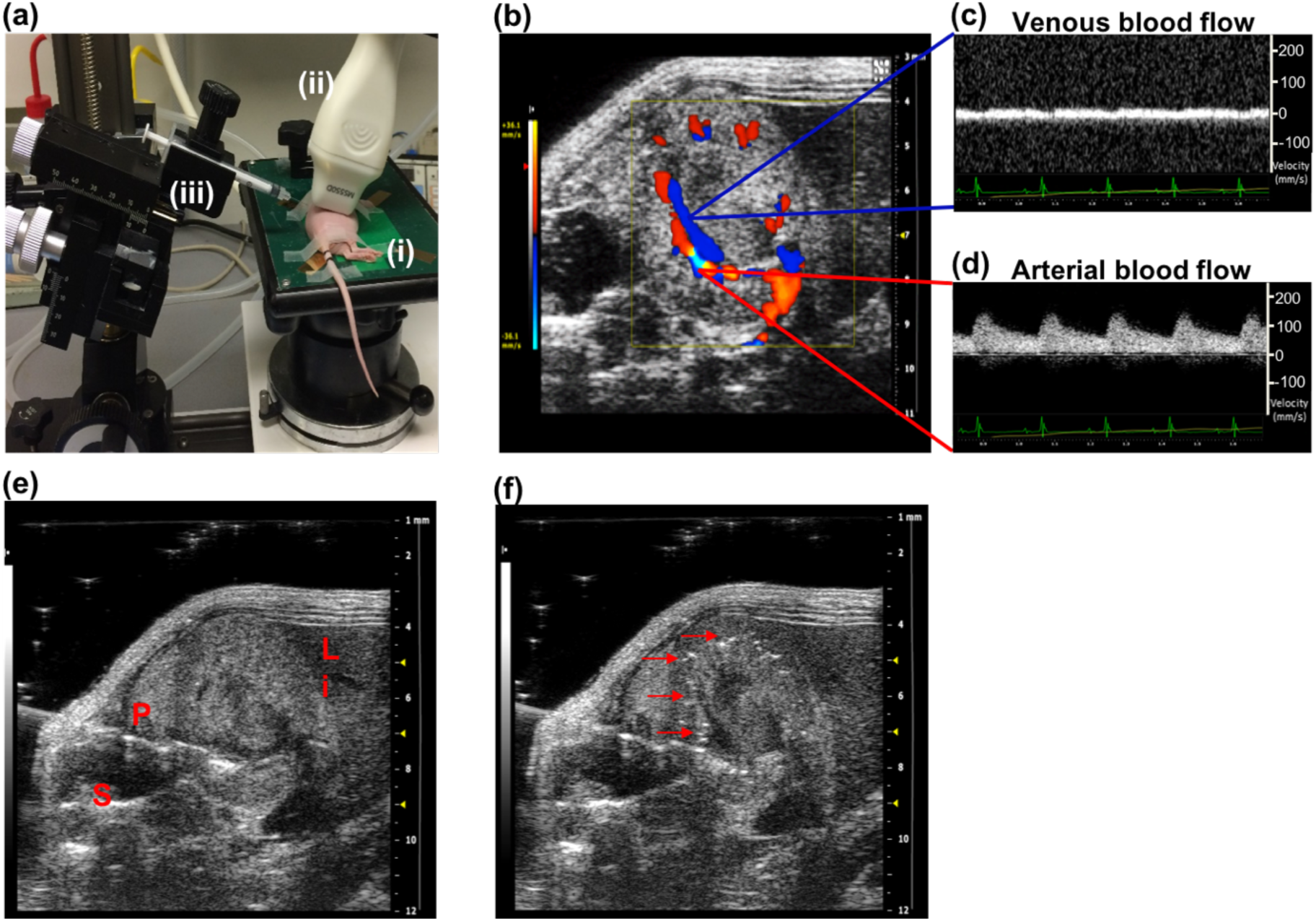
Non-invasive ultrasound-guided renal artery injection. (a) The animal was positioned in left lateral position on the platform (i) with the ultrasound transducer (ii) on the right lower quadrant of the abdomen and the syringe in injection mount (iii) facing towards the paravertebral muscle of the animal. (b) Colour Doppler image of the kidney showing renal blood supply (c & d) Pulsed wave Doppler images identifying the venous and arterial flow velocity pattern. (e) The needle was pierced through the skin and the vertebral muscle of the animal to penetrate into the right renal artery (labels: P = paravertebral muscle, S = spine, Li = liver). (f) The successful injection was visualised by the presence of hyperechogenic contrast around renal cortex (indicated by arrows).

### Assessment of the cell viability and cell localisation following renal artery injection

To assess the viability and localisation of GNR labelled ADSCs within the kidney, the mice were imaged with BLI and photoacoustic imaging serially for 7 days after ultrasound-guided right renal artery injection. *Ex vivo* BLI was performed at 1 hour and day 8 after injection to validate *in vivo* results.

At 1 hour after injection, BLI images showed the majority of BLI signal was coming from the right kidney (Figure **4a**, supported by *ex vivo* BLI, Figure **4d**) with some signal from the lungs. The percentage BLI signal in right kidney (relative to whole body BLI signal) was 29 ± 6 % (Figure **4b**). Immediately after BLI, PA images of both kidneys were acquired at 3-5 hours after injection (termed day 0) using a range of wavelength from 600 to 900 nm (Figure **4e** & **f**). PA images appeared to show a stronger PA signal within the right kidney compared to the left kidney at all wavelengths. However, images acquired at 730 nm showed the highest PA contrast in the right kidney than the surrounding renal blood vessels (Figure **4e**) consistent with the absorption peak of the GNRs (750 nm) which indicated the presence of GNR labelled cells within the right kidney. In addition, this signal was detected throughout the right kidney including cortex and medulla in 3D volume rendered PA images (Figure **4f**).

BLI images at days 1 and 3 after injection showed the persistence of the signal within the right kidney (Figure **4a**) and the percentage BLI signal was significantly higher than 1-hour post injection (P = 0.0063 & P = 0.0031, Figure **4b**). Similarly, PA images showed the signal intensity remained stable for 3 days (Figure **4g**). At day 7 after injection, BLI images showed a reduced signal within the right kidney (Figure **4a** supported by *ex vivo* BLI at day 8, Figure **4d**) and the percentage BLI signal was significantly lower than 1-hour post injection (P = 0.0093, Figure **4b**). However, high signal intensity was still detectable in PA images (Figure **4h**). Observation across the rest of the body showed the BLI signal intensity from the lungs and the whole body decreased over time (Figure **4c**) and no sign of cell migration to other organs was seen.

**Figure 4.**
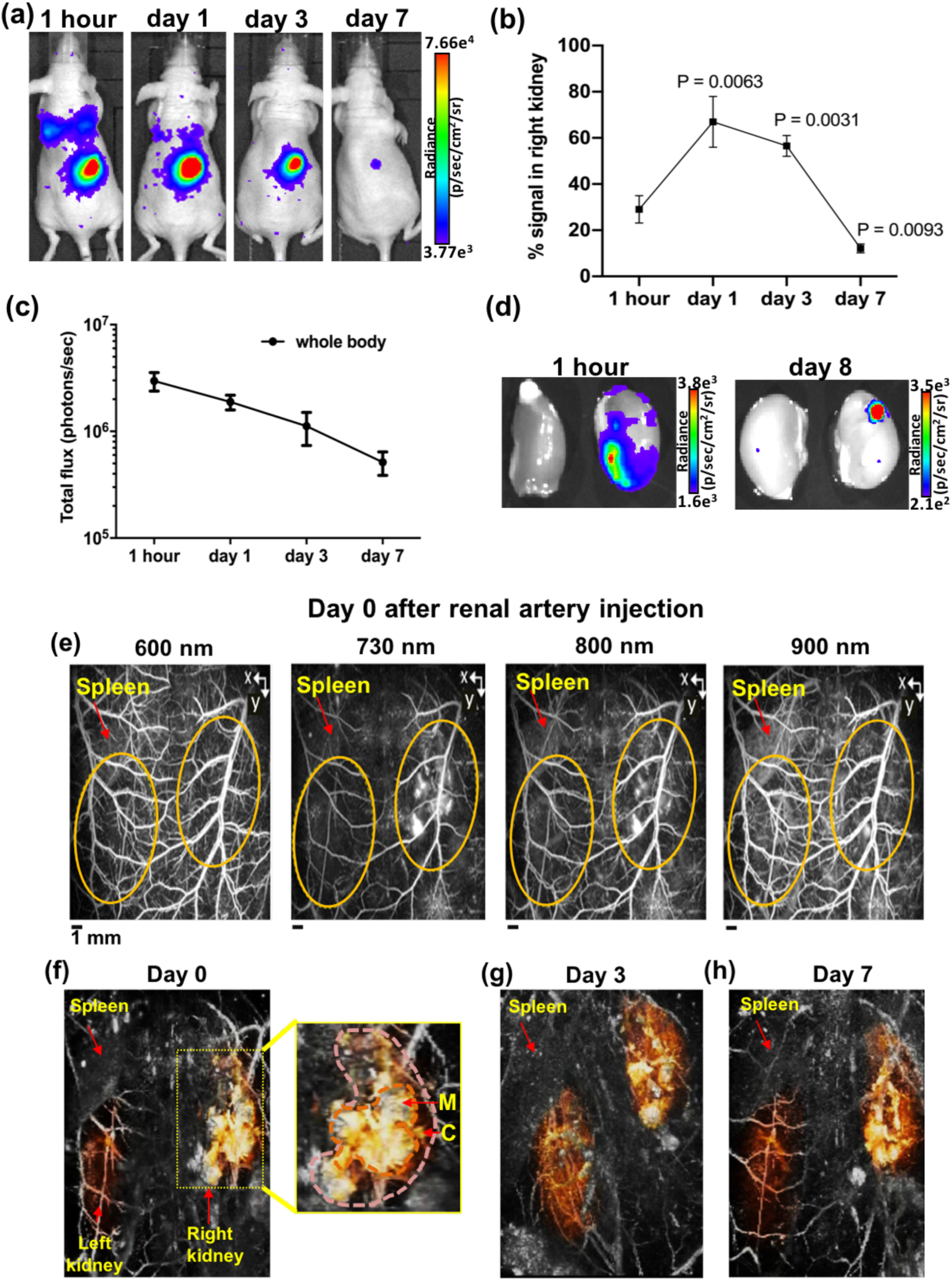
BLI and PA images of GNR labelled ADSC in right kidney at different time points after renal artery injection. (a) BLI images at 1 hour, days 1, 3 and 7 after injection showing the majority of signal within the right kidney and some signal from the lungs. (b) The percentage BLI signal in right kidney relative to whole body (1 hour vs days 1, 3 & 7). (c) BLI signal from the whole body decreased over time (photons/sec in log scale). (d) *Ex vivo* BLI images of both kidneys at 1 hour and day 8 after injection showing BLI signal from the right kidney. (e) PA images acquired at different wavelength showing the higher PA contrast in right kidney (indicated by yellow dotted lines) than the surrounding renal blood vessels at 730 nm using the spleen as anatomical landmark (images are presented as MIP). (f-h) 3D volume-rendered PA images at days 0, 3 and 7 after injection showing the widespread PA contrast in right kidney including cortex (C) and medulla (M), compared to left kidney (the kidneys are manually segment and false coloured). Data are shown as mean ± SD, n = 3.

### Histological analysis

In order to validate the presence of GNR labelled ADSCs in the right kidney after renal artery injection, the kidney sections from day 8 after injection were stained with human specific vimentin (ADSCs) and sliver enhancement (GNRs) using the left kidney as control tissue sections (Figure **5a**). Histological images showed widespread vimentin staining throughout the right kidney including cortex and medulla confirming the presence of ADSCs in the right kidney (Figure **5b**). Higher magnification images showed that areas positive for vimentin staining were also positive for sliver enhancement staining (Figure **5c** & **d**), confirming the retention of GNRs within ADSCs. In addition, the dual labelled cells were detected within the renal capillary bed (Figure **5c**) and renal parenchyma (Figure **5d**) compared to the left kidney tissue section. Observation of the pathology of right kidney showed no signs of trauma or haemorrhage resulting from the injection and no malignant tumour formation that could potentially result from ADSC engraftment.

**Figure 5.**
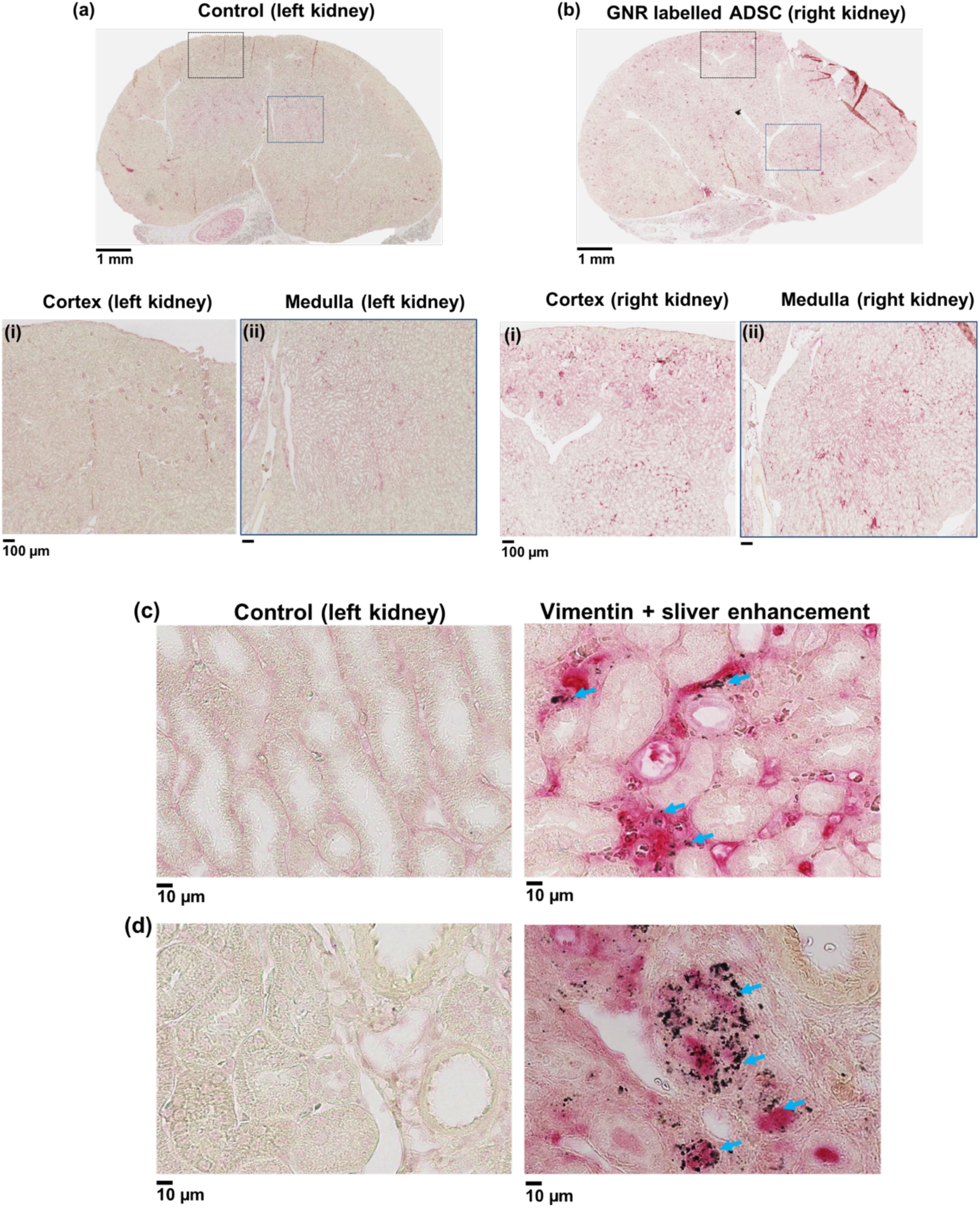
Histological analysis of GNR labelled ADSCs uptake in kidney at day 8 after right renal artery injection. (a & b) Vimentin (ADSC) staining of control (left kidney) and injected right kidney sections showing the widespeard red staining in the cortex and medulla of right kidney compared to the control. Positive vimentin (ADSC) and sliver enhancement staining of right kidney tissue sections showing the presence of dual labelled cells within the renal (c) capillary bed and (d) parenchyma compared to control (indicated by arrows, scale bar = 10 µm).

## Discussion

The aims of this study was to improve stem cell delivery to preclinical kidney models by developing a minimally invasive ultrasound-guided renal artery injection and to subsequently assess the viability and localisation of delivered cells by serial imaging by BLI and PA over seven days.

The intracellular uptake of GNRs by ADSCs was studied and the results showed that GNRs were internalised inside the cytoplasm. These results are consistent with studies carried out by *Comenge et al., 2016* who showed the entrapment of the same formulation of GNRs within the cytoplasmic organelles of MSCs^22^. The results from *in vitro* luciferase and differentiation assays demonstrated that GNRs labelling had no effect on ADSC proliferation and tri-lineage differentiation potential indicating that GNRs have no cytotoxic effect on ADSCs which is also consistent with the findings from *Comenge et al., 2016*.

Ultrasound-guided renal artery injection of GNR labelled cells demonstrated that this method is feasible, fast and less invasive than the routine renal artery injection using open abdomen surgical techniques. The main strength of the ultrasound-guided technique is a faster recovery and healing time with fewer complications such as infections in immunocompromised mice. In addition, the high temporal resolution of ultrasound allows for the optimisation of the injection techniques. During initial studies, the spillage of cells into the abdominal cavity was observed when the cells were re-suspended in PBS (results not shown). Since the alginate solution is known to be viscus^27^ and commonly used in ultrasound contrast agents^28^, the re-suspension medium was changed to 0.6% alginate solution to prevent spillage and to improve the accuracy of the injection by taking advantage of the ultrasound contrast provided by alginate.

The distribution of GNR labelled cells in the kidney after injection was assessed by using BLI and PA imaging. At 1 hour after injection, although the majority of the signal (29 ± 6%) was detected in the injected right kidney, there was some signal from the lungs in BLI images. This is most likely due to leakage of cells into the renal vein which is situated alongside the renal artery during the injection. Therefore, it is likely that the cells are transported to the lungs via venous drainage from the right ventricle as the blood from the renal vein drains into the inferior vena cava and then into the right atrium. Interestingly, BLI signal from the lungs decreased from day 1 after injection and was not detectable at day 7 while the percentage BLI signal in right kidney significantly increased at days 1 and 3 after injection. In comparison, many studies have reported the persistence of BLI signal within the lungs for 7 days after IV injection^7,29^. This is likely due to differences in the number of cells entrapped in the lung capillaries compared to the kidney.

More importantly, both BLI and PA images at the early time points suggested that the ultrasound-guided renal artery injection was successful and that the majority of the GNR labelled cells were retained within the right kidney and remained viable for 3 days. In addition, the widespread cell distribution across the kidney, including the cortex and medulla was detected in PA images and further confirmed by histological analysis. These results highlight the advantages of using PA imaging which can provide the high spatial resolution of cell localisation in the kidney.

Although consistent findings were seen between BLI and PA images at early time points, discrepancies were observed at day 7 after injection. Since BLI is a reporter gene-based cell tracking method that requires ATP, the signal generation depends on cell survival, while in PA imaging, the presence of signal only represents the existence of the direct labelling agent, GNR, not the cell itself. Therefore, if GNRs remain within the cell after cell death then the PA signal remains even when BLI decreases. These results consequently suggest that only a small number of cells remain viable 7 days after injection but the majority of dead cells are retained within the kidney. Nonetheless, these studies were done in mice with no underlying kidney disease and therefore the length of cell viability could be altered in disease state.

PA images were taken at different wavelengths to discern signal from the vasculature (600 nm) to that of GNR labelled cells (730 nm). Although the highest PA signal within the right kidney was at 730 nm without spectral unmixing it is unclear if some underlying signal came from haemorrhage. However histological findings at day 8 after injection confirmed the widespread presence of GNR labelled ADSCs throughout the kidney with no evidence of haemorrhage or injury derived from the injection of ADSCs and mice showed no evidence of blood within the urine at any point. Although histology also showed no signs of tumour formation there are still safety concerns regarding MSC-based therapies in kidney diseases. In addition, the maldifferentiation of MSC into glomerular adipocytes and sclerosis after renal artery injection have been reported in a rat model of renal failure^30^. Therefore, more studies are required to investigate the long term fate of MSCs in recipient’s organ after transplantation.

In summary, the results presented in this study demonstrated the successful development of a minimally invasive ultrasound-guided renal artery injection which can improve stem cell delivery to the kidney. In addition, the findings from this study displayed the benefits of combining a reporter gene (luciferase, BLI) with a direct cell labelling method (GNR, PA) which can provide both cell viability and cell localisation in the target organ. The new techniques which have been developed in this study can be applied to track other stem cell types such as kidney derived stem or progenitor cells. The information acquired from this study will can therefore be applied to the optimisation of cell therapies in various rodent models of kidney diseases such as acute kidney injury.

## Materials and methods

All chemical materials were from Sigma-Aldrich (Dorset, UK), unless otherwise stated.

### Gold nanorods (GNRs)

All experiments were performed using silica coated GNRs which were kindly provided by Dr Joan Comenge, Institute of Integrative Biology, University of Liverpool. The core size of GNR was 21 ± 3 nm in width and 49 ± 5 nm in length with silica shell thickness of 34 ± 2 nm. The absorption peak of GNR was 756 nm with the optical density of 0.74. Detailed synthesis and characterisation of GNRs were published in *Comenge et al., 2016*.

### Cell culture

All experiments were performed using human adipose derived stem cells (ADSCs) which were kindly provided by Dr Michelle Griffin, UCL Plastic and Reconstructive Surgery Department. ADSCs were transduced with a lentiviral vector using plasmid pSEW^31^ to express green fluorescence protein (GFP) and firefly luciferase^32^ under the control of the Friend murine leukaemia virus FB29 promoter. The successful transduction was confirmed by GFP expression using a fluorescence microscope (EVOS FL Auto cell imaging system, ThermoFisher Scientific, Massachusetts, USA) and the transduction efficiency was 42.56 ± 7.43%. The transduced ADSCs were grown in T175 flasks (Fisher Scientific, Loughborough, UK) in DMEM-F12, supplemented with 10% fetal calf serum (FCS, Invitrogen, Paisley, UK) in a humidified incubator at 37°C with 95% air and 5% CO_2_. Cells were grown to 80% confluence before being trypsinised, centrifuged for pelleting at 300 g, counted and then plated for *in vitro* experiments or GNR labelling.

### ADSC labelling with GNRs

3 × 10^5^ ADSCs were seeded in a T25 flask with 5 ml of DMEM-F12 supplemented with 10 % FCS and left to attach overnight. The next day, cells were incubated with GNRs at a concentration of 0.04235 mgAu/ml (80% medium + 20% GNRs in water) for 24 hours^22^. After washing 3 times with PBS, the labelled ADSCs were trypsinised, centrifuged for pelleting at 300 g, counted and then plated for in vitro studies or re-suspended in 0.6% alginate solution for *in vivo* cell injection.

### Silver enhancement staining

To visualise intracellular GNR uptake by ADSCs, silver enhancement staining was performed according to manufacturer’s instructions (silver enhancer kit, Sigma-Aldrich, St Louis, USA). GNR labelled ADSCs were plated in 24-well plates at a concentration of 0.22 × 10^5^ per well in triplicates and left to attach overnight. Next, the well plates were washed 3 times with PBS and the cells were fixed in 4% PFA for 30 minutes at room temperature. The cells were then washed 2 times with PBS followed by staining with silver enhancer mixture for 10 minutes at 20 °C. The cells were then washed 2 times with distilled water and fixed with 2.5% sodium thiosulfate solution for 3 minutes. Fixative was removed by washing 3 times with distilled water and the cells were counterstained with 1% Nuclear Fast Red and then imaged with EVOS FL Auto cell imaging system.

### *In vitro* luciferase assay

For luciferase-based cell proliferation assay, control ADSCs and GNR labelled ADSCs were plated in 96-well plates at a concentration of 5 × 10^3^ per well in triplicates. In vitro luciferase assay was performed using a Varioskan LUX multimode microplate reader (ThermoFisher Scientific, Waltham, MA USA) at days 1, 2, 3, 6 and 9 after plating. The measurements were acquired immediately after adding 300 μg/ml of D-luciferin and the results were presented as luminescence.

### Differentiation assay

To determine the effect of labelling on the differentiation potential of ADSCs, a differentiation assay was performed as previously described^33^. Control and GNR labelled ADSCs were plated in 24-well plates (Corning) at a concentration of 0.22 × 10^5^ per well in triplicates. When both control and labelled cells reached 90-100% confluency, the regular media was removed and differentiation media (adipogenic, chondrogenic or osteogenic) was added to appropriate wells and was changed every three days. Undifferentiated wells received regular media. After 3 weeks staining was conducted for each differentiation and the cells were imaged using the EVOS FL Auto cell imaging system.

### Adipogenic differentiation

Adipogenic differentiation was induced with DMEM-F12 medium containing 10% FCS, 1% penicillin/streptomycin, 10 ng/ml insulin, 1 μM dexamethasone, 500 µM 3-isobutyl-1-methylxanthine and 1 mM rosiglitazone. After 3 weeks, cells were fixed in 10% formalin for 30 minutes, washed with deionised water and then washed again with 60% isopropanol for 5 minutes prior to staining with Oil Red O working solution for 10 minutes. After staining, cells were washed several times with tap water.

### Chondrogenic differentiation

Chondrogenic differentiation was induced with DMEM-F12 medium containing 10% FCS, 1% penicillin/streptomycin, 0.1 μM dexamethasone, 50 μg/ml ascorbate, 10 ng/ml transforming growth factor (TGF) β1 (Life technologies, Paisley, UK) and 10 ng/ml insulin, transferrin, selenium. After 3 weeks, cells were fixed in 4% paraformaldehyde (PFA) for 30 minutes and then washed with deionised water. Then rinsed with 0.1 M HCl for 5 minutes and stained with Alcian Blue staining (1% in 0.1 M HCl) for 30 minutes. After staining the cells were washed with tap water 3 times.

### Osteogenic differentiation

Osteogenic differentiation was induced with DMEM-F12 medium containing 10% FCS, 1% penicillin/streptomycin, 0.1 μM dexamethasone, 10 mM β-glycerophosphate and 100 μg/ml ascorbate. After 3 weeks, cells were fixed in ice-cold 70% ethanol for 1 hour and washed with deionised water. Then the cells were stained with Alizarin Red staining (1% in deionised water, pH 4.1 - 4.3). After staining, cells were washed with tap water 3 times.

### *In vivo* studies

All animal studies were approved by the University College London Biological Services Ethical Review Committee and licensed under the UK Home Office regulations and the Guidance for the Operation of Animals (Scientific Procedures) Act 1986 (Home Office, London, United Kingdom). All mice were 6-8-week-old female CD-1 nude mice (Charles River Laboratories, UK) and all *in vivo* imaging experiments were performed under isoflurane anaesthesia (1.5% – 2.5% isoflurane in oxygen 1.5 - 2 l/min). The signs of injury resulting from injection and the general wellbeing of the animals were observed throughout the study by monitoring body weight and colour of urine.

### *In vivo* cell injections

Mice were imaged and injected using a VEVO 2100 ultrasound imaging system (VEVO 2100, FUJIFILM VisualSonics, Canada) with a Visualsonics ultrasound platform and injection rig. Mice were secured in the left lateral position with tape with their heart rate, body temperature and respiration rate monitored throughout the procedure. 2 × 10^5^ GNR labelled ADSCs (in 100 μl of 0.6 % alginate solution) were injected into the right renal artery of mice (n = 3) using ultrasound guidance. Total ultrasound injection and imaging took approximately 15 minutes, mice were then imaged serially with BLI and PA for 7 days.

### *In vivo* imaging

#### BLI

*In vivo* BLI was performed at 1 hour, days 1, 3 and 7 after renal artery injection using an IVIS Lumina (PerkinElmer, USA). Mice were injected intraperitoneally with 75 mg/kg D-luciferin (Promega) in 200 µl of PBS. Sequential BLI images were acquired 5 minutes after luciferin injection using auto exposure time with 0.5 minutes delay between two consecutive acquisitions. A rectangular region of interest (ROI) was placed over the whole body on the first image and subsequently pasted over every new image acquired until all ROIs reach their maximum intensity. The total signal in the ROI was quantified as total flux (photons/s) by using Living Image software version 4.5 (PerkinElmer). To calculate the percentage of BLI signal in right kidney, a circular ROI was placed over the right kidney and the signal from the right kidney ROI was divided by the whole body signal and presented as % signal in right kidney. Representative images were presented using radiance (the number of photons per second that leave a square centimetre of tissue and radiate into a solid angle of one steradian (sr) = p/sec/cm^2^/sr) as colour scale by utilizing the same software.

#### PA imaging

Directly after BLI imaging, mice were anaesthetised and PA scans were acquired at 3-5 hour after injection (termed day 0), and then at days 3 and 7 using a planar PA scanner (built in the Department of Medical Physics and Biomedical Engineering, UCL) based on a Fabry-Pérot (FP) polymer film ultrasound sensor and a tunable oscillator laser system (Quanta Ray Pro-270/premiScan; Newport Spectra Physics/GWU Lasertechnik). The PA images of right and left kidneys were acquired as previous described in *Ogunldade et al., 2018*^*26*^. In brief, the animals were placed in a supine position and a small amount of ultrasound gel was applied to the lower back of the animals. Laser light pulses were excited onto tissue at 7 ns with a pulse repetition frequency of 50 Hz. Photoacoustic signals were produced and detected by the FP ultrasound sensor using four different wavelengths between 600-900 nm (600, 730, 800 and 900 nm) to differentiate between signal derived from haemoglobin and that from GNR (however, spectral unmixing was not performed in this instance). An integrated heater and thermal chamber with temperature set to 34 °C was used during the imaging to maintain core body temperature. The images were displayed as MIPs and 3D volume-rendered images. False colouring, manual segmentation and extracting of the vasculature architecture was performed using Amira (FEI Visualization Sciences).

#### *Ex vivo* BLI imaging

*Ex vivo* BLI was performed at 1 hour and day 8 after renal artery injection. Under anaesthesia, mice were injected intraperitoneally with 75 mg/kg D-luciferin. At 10 minutes after injection, mice were sacrificed and the organs of interest were excised quickly and images were acquired using 5 minutes exposure time and binning 8.

#### Histology and immunohistochemistry (IHC)

At day 8 after renal artery injection, right and left kidneys were excised, fixed in 10% neutral buffered formalin and embedded in paraffin. Human specific vimentin (CONFIRM™ anti-Vimentin (V9) primary mouse monoclonal antibody, Roche) staining was performed on 5 μm kidney tissue sections to detect the presence of ADSCs using Ventana Discovery XT instrument and Ventana Red detection kit. For pre-treatment, Ventana CC1, equivalent to EDTA buffer, was used. To demonstrate the presence of GNRs, IHC stained sections were stained with silver enhancement staining (silver enhancer kit, Sigma-Aldrich, St Louis, USA) according to manufacturer’s instructions.

#### Statistical analysis

All *in vitro* experiments were repeated at least 3 times with 3 triplicates.

Statistical analysis was performed with GraphPad Prism version 6.01. Data were presented as mean ± standard deviation (SD). Unpaired two-tailed t test was conducted for the assessment of statistical significance.

## Acknowledgement

The authors would like to thank the EPSRC for the award of an Early Career Fellowship EP/L006472/1 (T.L.K, M.Z.T) and the MRC for funding via the UK Regenerative Medicine Platform Safety and Efficacy Hub MR/K026739/1 (P.S.P). D.J.S is supported by his British Heart Foundation intermediate basic science research fellowship (FS/15/33/31608) the BHF Centre for Regenerative Medicine (RM/17/1/33377) and an MRC Project Grant (MRC/PG/177291). We thank Dr. Sandra Meyer (FUJIFILM, VisualSonics) for expert advice on ultrasound-guided injection.

## Author contributions

M.Z.T, and T.L.K conceived the experiments and wrote the manuscript, M.Z.T, O.O., and D.J.S. conducted the experiments, P.S.P. performed lentiviral transduction, J.C. and R.L. synthesized gold nanorods, A.L.D, M.F.L and P.B assisted in writing manuscript.

## References

1. Imberti, B. et al. Insulin-like growth factor-1 sustains stem cell mediated renal repair. J Am Soc Nephrol 18, 2921–2928, doi:10.1681/asn.2006121318 (2007)

2. Yuan, L. et al. VEGF-modified human embryonic mesenchymal stem cell implantation enhances protection against cisplatin-induced acute kidney injury. Am J Physiol Renal Physiol 300, F207–218, doi:10.1152/ajprenal.00073.2010 (2011)

3. Si, X., Liu, X., Li, J. & Wu, X. Transforming growth factor-beta1 promotes homing of bone marrow mesenchymal stem cells in renal ischemia-reperfusion injury. International journal of clinical and experimental pathology 8, 12368–12378 (2015)

4. Peired, A. J., Sisti, A. & Romagnani, P. Mesenchymal Stem Cell-Based Therapy for Kidney Disease: A Review of Clinical Evidence. Stem Cells Int 2016, 4798639, doi:10.1155/2016/4798639 (2016)

5. Fischer, U. M. et al. Pulmonary passage is a major obstacle for intravenous stem cell delivery: the pulmonary first-pass effect. Stem Cells Dev 18, 683–692, doi:10.1089/scd.2008.0253 (2009)

6. Gooch, A. et al. Initial report on a phase I clinical trial: Prevention and treatment of post-operative Acute Kidney Injury with allogeneic Mesenchymal Stem Cells in patients who require on-pump cardiac surgery. Vol. 2 (2008)

7. Togel, F., Yang, Y., Zhang, P., Hu, Z. & Westenfelder, C. Bioluminescence imaging to monitor the in vivo distribution of administered mesenchymal stem cells in acute kidney injury. Am J Physiol Renal Physiol 295, F315–321, doi:10.1152/ajprenal.00098.2008 (2008)

8. Furlani, D. et al. Is the intravascular administration of mesenchymal stem cells safe? Mesenchymal stem cells and intravital microscopy. Microvascular research 77, 370–376, doi:10.1016/j.mvr.2009.02.001 (2009)

9. Alfarano, C. et al. Intraparenchymal injection of bone marrow mesenchymal stem cells reduces kidney fibrosis after ischemia-reperfusion in cyclosporine-immunosuppressed rats. Cell Transplant 21, 2009–2019, doi:10.3727/096368912x640448 (2012)

10. Bos, C. et al. In vivo MR imaging of intravascularly injected magnetically labeled mesenchymal stem cells in rat kidney and liver. Radiology 233, 781–789, doi:10.1148/radiol.2333031714 (2004)

11. Cai, J. et al. Maximum efficacy of mesenchymal stem cells in rat model of renal ischemia-reperfusion injury: renal artery administration with optimal numbers. PLoS One 9, e92347, doi:10.1371/journal.pone.0092347 (2014)

12. Liu, H. et al. The role of SDF-1-CXCR4/CXCR7 axis in the therapeutic effects of hypoxia-preconditioned mesenchymal stem cells for renal ischemia/reperfusion injury. PLoS One 7, e34608, doi:10.1371/journal.pone.0034608 (2012)

13. Hauger, O. et al. MR evaluation of the glomerular homing of magnetically labeled mesenchymal stem cells in a rat model of nephropathy. Radiology 238, 200–210, doi:10.1148/radiol.2381041668 (2006)

14. Sun, J. H., Teng, G. J., Ma, Z. L. & Ju, S. H. In vivo monitoring of magnetically labeled mesenchymal stem cells administered intravascularly in rat acute renal failure. Swiss Med Wkly 138, 404–412, doi:2008/27/smw-12000 (2008)

15. Hueper, K. et al. Acute kidney injury: arterial spin labeling to monitor renal perfusion impairment in mice-comparison with histopathologic results and renal function. Radiology 270, 117–124, doi:10.1148/radiol.13130367 (2014)

16. Hu, J. et al. Mesenchymal stem cells attenuate ischemic acute kidney injury by inducing regulatory T cells through splenocyte interactions. Kidney International 84, 521–531, doi:10.1038/ki.2013.114 (2013)

17. Grange, C. et al. Protective effect and localization by optical imaging of human renal CD133+ progenitor cells in an acute kidney injury model. Physiological reports 2, e12009, doi:10.14814/phy2.12009 (2014)

18. Jathoul, A. P. et al. Deep in vivo photoacoustic imaging of mammalian tissues using a tyrosinase-based genetic reporter. Nature Photonics 9, 239, doi:10.1038/nphoton.2015.22 https://www.nature.com/articles/nphoton.2015.22#supplementary-information (2015).

19. Urabe, K. et al. The inherent cytotoxicity of melanin precursors: a revision. Biochim Biophys Acta 1221, 272–278 (1994)

20. Lewinski, N., Colvin, V. & Drezek, R. Cytotoxicity of Nanoparticles. Small 4, 26–49, doi:10.1002/smll.200700595 (2008)

21. Li, W. & Chen, X. Gold nanoparticles for photoacoustic imaging. Nanomedicine (London, England) 10, 299–320, doi:10.2217/nnm.14.169 (2015)

22. Comenge, J. et al. Preventing Plasmon Coupling between Gold Nanorods Improves the Sensitivity of Photoacoustic Detection of Labeled Stem Cells in Vivo. ACS Nano 10, 7106–7116, doi:10.1021/acsnano.6b03246 (2016)

23. Berndl, E. S. L., He, X., Yuen, D. A. & Kolios, M. C. Photoacoustic imaging for assessing ischemic kidney damage in vivo. Vol. 10494 PWB (SPIE, 2018).

24. Pan, W. et al. Non-invasive detection of the early phase of kidney injury by photoacoustic/computed tomography imaging. Nanotechnology 29, 265101, doi:10.1088/1361-6528/aabcee (2018)

25. Scarfe, L. et al. Measures of kidney function by minimally invasive techniques correlate with histological glomerular damage in SCID mice with adriamycin-induced nephropathy. Scientific reports 5, 13601, doi:10.1038/srep13601 (2015)

26. Ogunlade, O. et al. In vivo three-dimensional photoacoustic imaging of the renal vasculature in preclinical rodent models. Am J Physiol Renal Physiol 314, F1145–f1153, doi:10.1152/ajprenal.00337.2017 (2018)

27. Lee, K. Y. & Mooney, D. J. Alginate: properties and biomedical applications. Progress in polymer science 37, 106–126, doi:10.1016/j.progpolymsci.2011.06.003 (2012)

28. Wheatley, M. A., Schrope, B. & Shen, P. Contrast agents for diagnostic ultrasound: development and evaluation of polymer-coated microbubbles. Biomaterials 11, 713–717 (1990)

29. Cao, J. et al. In Vivo Tracking of Systemically Administered Allogeneic Bone Marrow Mesenchymal Stem Cells in Normal Rats through Bioluminescence Imaging. Stem Cells International 2016, 3970942, doi:10.1155/2016/3970942 (2016)

30. Kunter, U. et al. Mesenchymal stem cells prevent progressive experimental renal failure but maldifferentiate into glomerular adipocytes. J Am Soc Nephrol 18, 1754–1764, doi:10.1681/asn.2007010044 (2007)

31. Demaison, C. et al. High-level transduction and gene expression in hematopoietic repopulating cells using a human immunodeficiency [correction of imunodeficiency] virus type 1-based lentiviral vector containing an internal spleen focus forming virus promoter. Human gene therapy 13, 803–813, doi:10.1089/10430340252898984 (2002)

32. Law, G. H., Gandelman, O. A., Tisi, L. C., Lowe, C. R. & Murray, J. A. Mutagenesis of solvent-exposed amino acids in Photinus pyralis luciferase improves thermostability and pH-tolerance. The Biochemical journal 397, 305–312, doi:10.1042/bj20051847 (2006)

33. New, S. E. et al. A matter of identity - Phenotype and differentiation potential of human somatic stem cells. Stem Cell Res 15, 1–13, doi:10.1016/j.scr.2015.04.003 (2015)

